# Sleep deprivation reduces the density of individual spine subtypes in a branch-specific fashion in CA1 neurons

**DOI:** 10.1101/2021.06.30.450529

**Authors:** Youri G Bolsius, Peter Meerlo, Martien J Kas, Ted Abel, Robbert Havekes

## Abstract

Sleep deprivation has a negative impact on hippocampus-dependent memory, which are thought to depend on cellular plasticity. We previously found that five hours of sleep deprivation robustly decreases dendritic spine density in the CA1 area of the hippocampus in adult male mice. However, recent work by others suggests that sleep deprivation increases the density of certain spine types on specific dendritic branches. Based on these recent findings and our previous work, we conducted a more in-depth analysis of different spine types on branches 1, 2 and 5 of both apical and basal dendrites to assess whether five hours of sleep deprivation may have previously unrecognized spine-type and branch-specific effects. This analysis shows no spine-type specific changes on branch 1 and 2 of apical dendrites after sleep deprivation. In contrast, sleep deprivation decreases the number of mushroom and branched spines on branch 5. Likewise, sleep deprivation reduces thin, mushroom, and filopodia spine density on branch 5 of the basal dendrites, without affecting spines on branch 1 and 2. Our findings indicate that sleep deprivation leads to local branch-specific reduction in the density of individual spine types, and that local effects might not reflect the overall impact of sleep deprivation on CA1 structural plasticity. Moreover, our analysis underscores that focusing on a subset of dendritic branches may lead to potential misinterpretation of the overall impact of in this case sleep deprivation on structural plasticity.

## 1. Introduction

A lack of sleep is a common problem in our 24/7 society, and has a detrimental impact on brain functioning. Work by numerous laboratories indicated that the hippocampus is one of the brain regions particularly vulnerable to sleep deprivation (reviewed in Havekes & Aton, 2020; Kreutzmann, Havekes, Abel, & Meerlo, 2015; Raven, Van der Zee, Meerlo, & Havekes, 2018). Sleep deprivation not only impacts fundamental hippocampus-dependent processes such as memory, but can also lead to the shrinkage of hippocampal volume (Novati, Hulshof, Koolhaas, Lucassen, & Meerlo, 2011).

Biochemical analyses of adult mice hippocampi revealed that as little as 5-6 hours of sleep deprivation leads to a numerous of changes in molecular and cellular processes (Havekes, Vecsey, & Abel, 2012) including the overall reduction of spine density in the hippocampal subregions CA1 (Acosta-peña et al., 2015; Havekes et al., 2016; Wong, Tann, Ibanez, & Sajikumar, 2019) and dentate gyrus (DG) (Raven, Meerlo, Zee, Abel, & Havekes, 2019), but not in area CA3 (Havekes et al., 2016). Further examination of the CA1 subregion revealed that the reduction in spine density is similar across apical and basal dendrites (Havekes et al., 2016; Wong et al., 2019). However, the reduction in spine density is not uniform along dendrites, but varies depending on the distance from soma and branch number (Havekes et al., 2016; Raven et al., 2019). Indeed, sleep deprivation decreases spine numbers in both apical and basal CA1 neurons at a distance of approximately 30-150 μm from the soma (Havekes et al., 2016), whereas in the DG, the reduction in spines are found closer to the soma (0-30 μm) (Raven et al., 2019). Furthermore, only specific branches (*i.e.* branch 3-8 of CA1 apical/basal dendrites and branch 1-4 of both blades in the DG) show significant decreases in spine numbers following sleep deprivation (Raven et al., 2019). Such branch-specific changes in response to learning and sleep have also been reported for the motor cortex (Yang et al., 2014). With respect to spine type, it should be noted that sleep deprivation leads to an overall reduction of all spine types in CA1 neurons, whereas it specifically affects thin and branched spines in the DG (Havekes et al., 2016; Raven et al., 2019). Together, these observations indicate that sleep deprivation may lead to local changes in structural plasticity that are specific for hippocampal subregion, dendritic branch, and spine type.

Despite the general consensus that sleep deprivation has a negative impact on hippocampal connectivity (Acosta-peña et al., 2015; Havekes et al., 2016; Raven et al., 2019; Wong et al., 2019) other studies reported opposing findings (De Vivo et al., 2017; Gisabella, Scammell, Bandaru, & Saper, 2020; Spano et al., 2019). Of special interest is the work from Gisabella and colleagues which uses comparable experimental parameters, like the sleep deprivation method and age of animals. They observed an overall increase in density of CA1 spines of sleep-deprived mice. Interestingly, the observed increase was found specifically for thin spines, and profoundly in distal (150-300 μm) regions from the soma of apical dendrites, while mushroom spines were exclusively increased in basal dendrites more close to the soma (0-150 μm) (Gisabella et al., 2020).

Prompted by these at a first glance opposing findings by Gisabella and colleagues (Gisabella et al., 2020), we decided to conduct a further in-depth analysis of our previously gathered spine data (Havekes et al., 2016). This additional analysis examined the impact of sleep deprivation on different spine subtypes at a branch-specific level. Because Gisabella et al (2020) restricted their study to the first two branches stemming from the main apical or basal dendrite, we determined the density of specific spine subtypes along the first and second branches of both apical and basal dendrites. We will also include the fifth branch as a representative for branches 3-8, which were all similarly affected by sleep deprivation (Havekes et al., 2016). Altogether our current study aims to elucidate the origin of the seemingly contradictory findings with respect to the impact of sleep deprivation on structural plasticity in CA1 neurons.

## 2. Materials and Methods

For this study, we re-analyzed previously gathered and published data from Havekes et al (2016) and briefly summarized the experimental procedures below. Three-months old male C57BL/6J mice were obtained from Jackson Laboratories. Upon arrival, mice were grouped housed with four littermates on a 12h:12h light:dark cycle (lights on at 7 am) with food and water available *ad libitum*. One week prior to the start of the experiment all mice were single housed. Mice were sleep deprived for 5 hours at the start of the lights on period (7 am), using the gentle handling method (Havekes & Aton, 2020; Havekes et al., 2016; Raven et al., 2019). After sleep deprivation, both experimental groups were immediately sacrificed and the brains were impregnated using the Rapid Golgi stain kit (FD Neurotechnologies Inc., Columbia, MD, USA) and coronal 80μm thick hippocampal sections were prescreened to find neurons qualified for spine analysis. We used a stereology-based software (Neurolucida, v10, Microbrightfield, VT), and a Zeiss Axioplan 2 image microscope with an Optronics MicroFire CCD (1600 × 1200) digital camera, motorized in X, Y, and Z-focus for high-resolution image acquisition and digital quantification in combination with a 100x objective. Dorsal hippocampus slices, with anterior-posterior coordinates between −1.5 and −2.30, were prescreened for impregnated CA1 pyramidal neurons along the anterior/posterior axis to see if they were qualified for further analysis. Pyramidal CA1 neurons that were insufficiently impregnated or truncated were excluded for further analysis. After prescreening, five neurons per animal were selected for further spine analysis. For the visualization of the spines, we used a 100x Zeiss objective lens with immersion oil, to identify the spine subclass. Prescreening and analysis was done by an experimenter blind to treatment. See Havekes et al (2016) for a more detailed description regarding data collection. All experiments were conducted according to National Institutes of Health guidelines for animal care and were approved by the Institutional Animal Care and Use Committee of the University of Pennsylvania.

For the more in-depth analysis of the current study, we focused on apical and basal dendrites of the CA1 pyramidal neurons (Fig.1A), and analyzed the first, second, and fifth branch in which we investigated whether sleep deprivation affects the different subtypes of spines in a branch-specific fashion (Fig.1B-D). These specific dendritic branch numbers were deliberately chosen in such a way that it includes branches investigated previously in our work (Havekes et al., 2016) and more recently by others (Gisabella et al., 2020). Importantly, we used the centrifugal branch ordering method to label branches for spine counting (Fig.1B). This branch counting method starts from the soma, and new branch numbers are assigned after every burification (or trification) of the dendrite. Spines were analyzed using Neurolucida. The neuronal tracing was conducted by an experimenter blind to treatment at Neurodigitech (San Diego, CA, USA). We analyzed data from 5 neurons per animal and 5-6 animals per group. The chosen neurons were fully traced and the data from these 5-6 neurons per animal were averaged leading to a single data point per animal. Data sets were analyzed using non-paired t-tests with adjusted p-values for multiple testing according to the Benjamini-Hochberg method.

**Figure 1:**
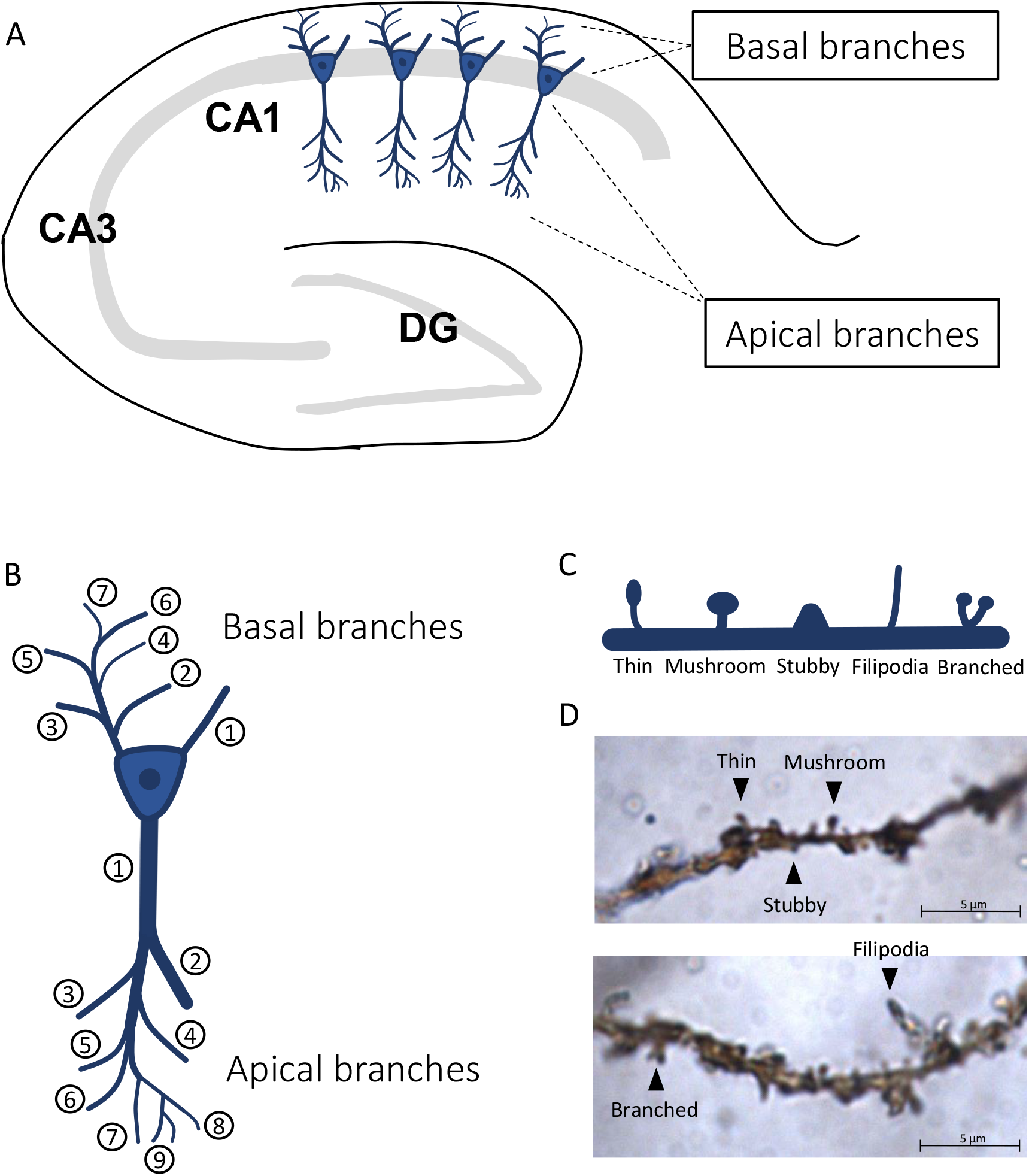
Overview of CA1 pyramidal neurons, branch numbering and spine subtypes. (A) A schematic representation of the hippocampus including the CA1 pyramidal neurons. (B) Illustration of the assignment of branch numbers to dendritic branches (centrifugal branch ordering method). (C) Overview of the different spine-subtypes analyzed in this study. (D) Example of golgi-stained pyramidal dendrites with different spine subtypes. The example pictures were taken from animals of the non-sleep deprivation group (NSD).

## 3. Results

### Sleep deprivation leads to branch-specific reduction in spines along apical dendrites

We previously reported that sleep deprivation causes an overall reduction in spine density selectively on apical branches 3 to 8 (Fig.2A; Havekes et al., 2016). Our new analysis on branch 1, 2, and 5 now indicates that the effects of sleep deprivation on spine density are branch specific. Particularly, after correcting the p-value for multiple testing, we found no effect of sleep deprivation on the number of spines selectively on branch 1 (Fig.2B), neither did sleep deprivation impact the number of dendritic spines for any subtype on branch 2 (Fig.2C). In contrast, sleep deprivation caused a decrease in the number of mushroom spines (p=0.004) and branched spines (p=0.045) on branch 5 (Fig.2D).

**Figure 2:**
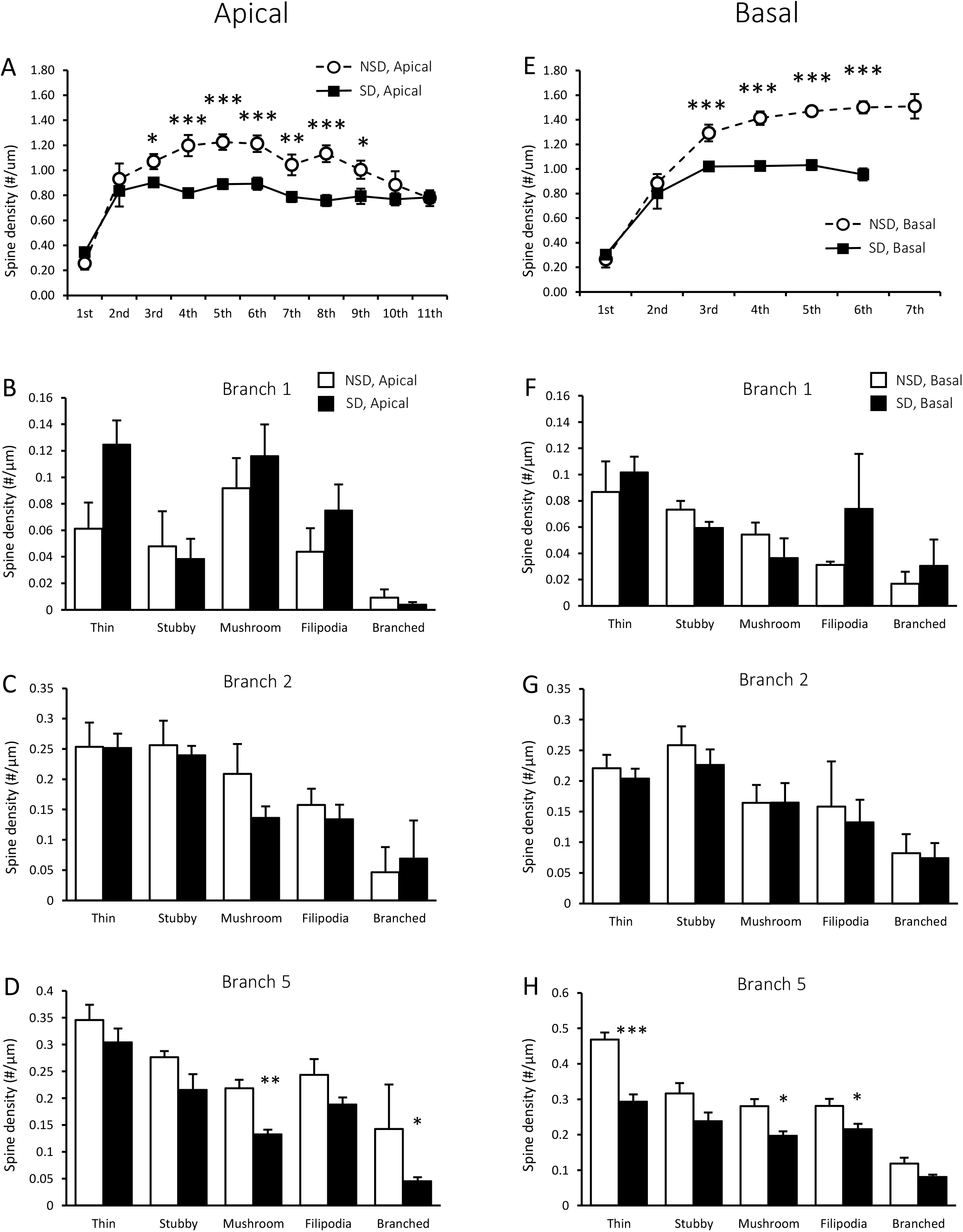
The effects of sleep deprivation on spine density in CA1 pyramidal neurons are spine-type and branch specific. (A) Sleep deprivation reduces the overall spine density on the apical dendritic branches 3 to 8 in CA1 (figure copied from supplementary fig.1H from Havekes et al., 2016 with minor graphic modifications). (B) For all spine types, no significant changes were observed in the number of spines on branch 1 (NSD, n=6, SD, n=5). (C) Sleep deprivation does not alter the density of any spine type on branch 2 (all groups, n=6), (D) while it does lead to a reduction of mushroom and branched spines on branch 5 (all groups, n=6). (E) Sleep deprivation reduces the overall spine density on the dendritic branches 3 to 6 on basal CA1 neurons (figure copied from supplementary fig.1G from Havekes et al., 2016). (F, G) Sleep deprivation does not alter the overall spine density on the first and second basal branches (all groups, n=6) (H) However, Sleep deprivation reduces the density of thin and mushroom spines, as well as the number of filipodia on branch 5 (all groups, n=6). Data are presented as mean plus s.e.m. *p <0.05, **p<0.005, ***p <0.001.

### Sleep deprivation leads to branch-specific reduction in spines along basal dendrites

Overall spine densities on branches 3 to 6 were significantly reduced by sleep deprivation. (Fig.2E; Havekes et al., 2016). Spine-specific analyses indicated that sleep deprivation did not alter the density of any type of spine for branch 1 and 2 (Fig.2F, G). In contrast, sleep deprivation attenuated the number of thin (p<0.001) and mushroom spines (p=0.011), as well as the number of filopodia (p=0.042) on branch 5 (Fig.2H). The observed changes in spine densities in both apical and basal dendrites largely returned to control levels after 3 hours of recovery sleep (no data shown).

## 4. Discussion

Our spine specific analysis revealed that sleep deprivation alters the density of individual spine types in a branch-specific fashion in CA1 neurons. Albeit the observed tendency towards an increase in the number of thin spines on apical branch 1, there was no significant increase in thin spines after correcting for multiple testing. Also on apical branch 2, the density of any spine type was not significantly altered after sleep loss. Furthermore, sleep deprivation decreases the number of mushroom and branched spines on the apical dendrites of branch 5 and reduces the density of thin, mushroom, and filopodia spines on the basal dendrites of branch 5. These findings indicate that, depending on the branch number examined, sleep deprivation reduces the number of specific spine subtypes on apical and basal dendrites. Hence, sleep deprivation has local effects in structural plasticity that might not reflect the overarching impact of sleep deprivation on CA1 spine density (*e.g*. Havekes et al., 2016).

The observed tendency towards increased thin spines exclusively along the first branch following sleep deprivation was not significant after correcting for multiple testing. This non-significant tendency is in line with the work of Gisabella et al (2020) suggesting that sleep deprivation might increase the number of thin spines in CA1 neurons. It is important to note that Gisabella et al (2020) used a different branch counting method (shaft ordering), and limited their analysis to the first two branches, leaving all other branches unanalyzed. As a result, it is unclear whether the changes they reported are representative for all branches or, alternatively, are branch-specific and not reflecting the overall impact of sleep deprivation. Indeed, our results suggest that local changes might not necessarily reflect the overarching negative effect of sleep deprivation on structural plasticity reported by us and others (*e.g*. Acosta-peña et al., 2015; Havekes & Aton, 2020; Havekes et al., 2016; Raven et al., 2019, 2018; Wong et al., 2019). As such, these findings together with our previous work underscore the importance of tracing all branch numbers when examining the impact of a manipulation on structural plasticity.

Our data also suggests that the discrepancy in literature concerning the impact of sleep deprivation on spines might not be related to the method of spine visualization, but is rather depending on the dendritic branch examined. Whereas Gisabella et al (2020) used a genetic and viral approach to visualize their spines, the current study uses a different approach, the Golgi-Cox staining method. Although both visualization approaches differ, it seems that the results are not contradictive and are even relatively comparable if similar branches are examined. Moreover, our previous study (Havekes et al., 2016) compared the changes in spines between Golgi stained and Dil stained CA1 neurons, revealing similar spine alterations following sleep deprivation. Nevertheless, it should be noted that Golgi only stains a small fraction of the CA1 pyramidal neurons. However, this seems not to influence the impact of sleep deprivation on the spine changes.

While previous studies found that a large number of molecular mechanisms are affected by sleep deprivation (*e.g*. Havekes et al., 2016), none of these identified molecular mechanisms can easily explain the observed branch- and spine type-specific alterations on the CA1 pyramidal neurons. Spine plasticity is determined by pre-synaptic input (or lack of input) and neuronal activity. A previous study by Yang et al (2014) demonstrated that sleep causes branch-specific spine formation after a learning trial due to enhanced reactivation of neurons involved in the learning process during non-REM sleep (Yang et al., 2014). Although our Golgi staining does not provide any information about the pre-synaptic input, it is known that the CA1 receives input from both area CA3 and cortical regions, and that some of these brain areas are showing aberrant activity patterns after sleep deprivation (Delorme, Kodoth, & Aton, 2019). Interestingly, proximal dendrites are more prone to receive inputs from closer sources such as the Schaffer collaterals of the CA3, whereas the input of distal dendrites are more related to cortical and thalamic inputs (Spruston, 2008). Moreover, CA1 innervation by interneurons can be highly branch specific, and can even target specific areas on individual dendritic branches (Bloss et al., 2016). A recent study found that the input of CA1 PV^+^ interneurons to pyramidal cells is altered during post-learning sleep deprivation (Ognjanovski, Broussard, Zochowski, & Aton, 2018). Altogether, it remains unclear how these specific and heterogeneous input pathways are affected by sleep deprivation. Future studies examining these specific inputs may shed light on the reported local changes in CA1 structural plasticity and its functional consequences.

## Acknowledgements

This work has been supported by NiH grant R01 MH117964 (T. Abel, PI)

## Notes

**Conflicts of interest:** None

### Competing Interest Statement

The authors have declared no competing interest.

## References

Acosta-peña, E., Camacho-Abrego, I., Melgarejo-Gutiérrez, M., Flores, G., Drucker-Colín, R., & García-García, F. (2015). Sleep deprivation induces differential morphological changes in the hippocampus and prefrontal cortex in young and old rats. Synapse, 69(1), 15–25. https://doi.org/10.1002/syn.21779

Bloss, E. B., Cembrowski, M. S., Karsh, B., Colonell, J., Fetter, R. D., & Spruston, N. (2016). Structured Dendritic Inhibition Supports Branch-Selective Integration in CA1 Pyramidal Cells. Neuron, 89(5), 1016–1030. https://doi.org/10.1016/j.neuron.2016.01.029

De Vivo, L., Bellesi, M., Marshall, W., Bushong, E. A., Ellisman, M. H., Tononi, G., & Cirelli, C. (2017). Ultrastructural evidence for synaptic scaling across the wake/sleep cycle. Science, 355(6324), 507–510. https://doi.org/10.1126/science.aah5982

Delorme, J. E., Kodoth, V., & Aton, S. J. (2019). Sleep loss disrupts Arc expression in dentate gyrus neurons. Neurobiology of Learning and Memory, 160(January 2018), 73–82. https://doi.org/10.1016/j.nlm.2018.04.006

Gisabella, B., Scammell, T., Bandaru, S. S., & Saper, C. B. (2020). Regulation of hippocampal dendritic spines following sleep deprivation. Journal of Comparative Neurology, 528(3), 380–388. https://doi.org/10.1002/cne.24764

Havekes, R., & Aton, S. J. (2020). Impacts of Sleep Loss versus Waking Experience on Brain Plasticity: Parallel or Orthogonal? Trends in Neurosciences, 43(6), 385–393. https://doi.org/10.1016/j.tins.2020.03.010

Havekes, R., Park, A. J., Tudor, J. C., Luczak, V. G., Hansen, R. T., Ferri, S. L., … Abel, T. (2016). Sleep deprivation causes memory deficits by negatively impacting neuronal connectivity in hippocampal area CA1. ELife, 1–22. https://doi.org/10.7554/eLife.13424

Havekes, R., Vecsey, C. G., & Abel, T. (2012). The impact of sleep deprivation on neuronal and glial signaling pathways important for memory and synaptic plasticity. Cellular Signalling, 24(6), 1251–1260. https://doi.org/10.1016/j.cellsig.2012.02.010

Kreutzmann, J. C., Havekes, R., Abel, T., & Meerlo, P. (2015). Sleep deprivation and hippocampal vulnerability: changes in neuronal plasticity, neurogenesis and cognitive function. Neuroscience, 309, 173–190. https://doi.org/10.1016/j.neuroscience.2015.04.053

Novati, A., Hulshof, H. J., Koolhaas, J. M., Lucassen, P. J., & Meerlo, P. (2011). Chronic sleep restriction causes a decrease in hippocampal volume in adolescent rats, which is not explained by changes in glucocorticoid levels or neurogenesis. Neuroscience, 190, 145–155. https://doi.org/10.1016/j.neuroscience.2011.06.027

Ognjanovski, N., Broussard, C., Zochowski, M., & Aton, S. J. (2018). Hippocampal network oscillations rescue memory consolidation deficits caused by sleep loss. Cerebral Cortex, 28(10), 3711–3723. https://doi.org/10.1093/cercor/bhy174

Raven, F., Meerlo, P., Zee, E. A. Van Der, Abel T., & Havekes, R. (2019). A brief period of sleep deprivation causes spine loss in the dentate gyrus of mice. Neurobiology of Learning and Memory, (160), 83–90.

Raven, F., Van der Zee, E. A., Meerlo, P., & Havekes, R. (2018). The role of sleep in regulating structural plasticity and synaptic strength: Implications for memory and cognitive function. Sleep Medicine Reviews, 39, 3–11. https://doi.org/10.1016/j.smrv.2017.05.002

Spano, G. M., Banningh, S. W., Marshall, W., de Vivo, L., Bellesi, M., Loschky, S. S., … Cirelli, C. (2019). Sleep Deprivation by Exposure to Novel Objects Increases Synapse Density and Axon-Spine Interface in the Hippocampal CA1 Region of Adolescent Mice. The Journal of Neuroscience□: The Official Journal of the Society for Neuroscience, 39(34), 6613–6625. https://doi.org/10.1523/JNEUROSCI.0380-19.2019

Spruston, N. (2008). Pyramidal neurons: Dendritic structure and synaptic integration. Nature Reviews Neuroscience, 9(3), 206–221. https://doi.org/10.1038/nrn2286

Wong, L.-W., Tann, J. Y., Ibanez, C. F., & Sajikumar, S. (2019). The p75 Neurotrophin Receptor Is an Essential Mediator of Impairments in Hippocampal-Dependent Associative Plasticity and Memory Induced by Sleep Deprivation. The Journal of Neuroscience, 39(28), 5452–5465. https://doi.org/10.1523/jneurosci.2876-18.2019

Yang, G., Lai, C. S. W., Cichon, J., Ma, L., Li, W., & Gan, W.-B. (2014). Sleep promotesbranch-specific formation ofdendritic spines after learning. Science, 344(6188), 1173–1178.

